# Privacy-Preserving Microbiome Analysis Using Secure Computation

**DOI:** 10.1101/025999

**Authors:** Justin Wagner, Joseph N. Paulson, Xiao-Shaun Wang, Bobby Bhattacharjee, Héctor Corrada Bravo

## Abstract

**Motivation:** Developing targeted therapeutics and identifying biomarkers relies on large amounts of patient data. Beyond human DNA, researchers now investigate the DNA of micro-organisms inhabiting the human body. An individual’s collection of microbial DNA consistently identifies that person and could be used to link a real-world identity to a sensitive attribute in a research dataset. Unfortunately, the current suite of DNA-specific privacy-preserving analysis tools does not meet the requirements for microbiome sequencing studies.

**Results:** We augment an existing categorization of genomic-privacy attacks to incorporate microbiome sequencing and provide an implementation of metagenomic analyses using secure computation. Our implementation allows researchers to perform analysis over combined data without revealing individual patient attributes. We implement three metagenomic analyses and perform an evaluation on real datasets for comparative analysis. We use our implementation to simulate sharing data between four policy-domains and measure the increase in significant discoveries. Additionally, we describe an application of our implementation to form patient pools of data to allow drug companies to query against and compensate patients for the analysis.

**Availability:** The software is freely available for download at: http://cbcb.umd.edu/∼hcorrada/projects/secureseq.html

## 1 Introduction

A significant application of DNA sequencing is studying the microbial communities that inhabit the human body. It is estimated that only 1 in 10 cells that reside in and on a person contain that individual’s DNA [22]. Microbiome sequencing seeks to characterize and classify all of these non-human cells. Most of these microbes cannot be cultured and studied in the laboratory, therefore direct sequencing is used. Of current interest in metagenomics is determining the relationship between microbiome features and identifying disease-causing bacteria.

The Human Microbiome Project [23], the Global Enterics Multi-Center Study [20], the Personal Genome Project [5], and the American Gut Project [4] aim to characterize the ecology of human microbiota and the impacts on human health. Potentially pathogenic or probiotic bacteria can be identified by detecting significant differences in their distribution across healthy and disease populations. While the biology has led to promising results, the privacy implications of microbiome analysis are just now being identified with no secure analysis tools available.

We review recent work showing how metagenomics data is a unique identifier across datasets and could be used to link an attribute to an individual [8]. To counter these concerns, we present an implementation and evaluation of metagenomic association analyses in a secure multiparty computation (SMC) framework. For our work, we use garbled circuits, a technique from cryptography for private computation between two parties. We provide a detailed review of this approach in Section 3.

We believe that implementing metagenomic analyses in an SMC framework will prove beneficial to the secure computation community as well as researchers focused on the human microbiome. Computational biologists will benefit from a method that allows quick and private computation over datasets which they may be obligated to keep confidential. Security researchers can draw on the findings from our work and construct protocols that enable sharing large, sparse datasets to perform analysis. Additionally, this work provides a mechanism for groups of patients to construct and manage high-quality reference datasets.

In summary, we detail the following contributions in this work:

- An analysis of microbiome-privacy concerns by expanding a categorization of genomic-privacy attacks. This provides a systematization of privacy concerns for metagenomic analyses and acts as a resource for researchers to address domain-specific problems.
- A secure computation implementation of widely-used microbiome analyses. We benchmark the implementation on three datasets used by metagenomic researchers. We also quantify the statistical gain a researcher will experience from using our tool by simulation with a dataset that contains samples from four different countries.

## 2 Problem Overview

In this section we describe the privacy threats of microbiome data and annotate them according to an existing categorization of genome privacy risks. We provide a comprehensive review of microbiome sequencing and metagenomics in the Supplementary Note, Section 1.

### 2.1 Forensic Identification

One prominent study proved that a person’s hand bacteria can identify objects that individual touched [7]. The authors first show the bacteria left after touching a keyboard are separate and unique between individuals. To measure the stability of the bacterial community left behind on the keyboards, the authors compared sequencing results for keyboard samples from the same person stored for 3 to 14 days at -20 degrees C and room temperature. The community makeup for each sample was not significantly different between any sample storage method. Next the authors calculated the UniFrac distance in community membership between keyboard samples from nine people and a database of microbiome samples from 270 individual’s hands. The closest match for each sample was the individual who touched the keyboard. This study was the first to show the identification power of an individual’s microbiome signature.

### 2.2 Identification with Metagenomic Codes

A recent analysis showed that metagenomic data alone can uniquely identify individuals in the Human Microbiome Project dataset [8]. The authors build minimal hitting sets to find a collection of microbiome features that are unique to each individual compared to all others in a dataset. The minimal hitting set algorithm was built using four types of features - OTUs, species, genetic markers, and thousand base windows matching reference genomes. The authors use a greedy algorithm and prioritize features by abundance gap, the difference in abundance between a feature in one sample compared to all other samples. The authors called these sets of features ”metagenomic codes” and used the codes built at the first time point in the Human Microbiome Project dataset to match individuals at a second time point. The genetic marker and base window codes were the best identifiers between the two time points. The OTU and species level codes also identified individuals but had a higher false-positive rate. As the authors note, the discovery of an identifiable microbiome fingerprint substantially changes the considerations for publicly releasing human microbiome data.

### 2.3 Genetic Re-identification Attacks

Through detailing attacks on genetic datasets, a recent article provided a categorization of techniques to breach participant privacy [6]. The attacks fall into several areas: *Identity Tracing* defined as determining the identity of an anonymized DNA sample using non-private attributes, *Attribute Disclosure* which uses a piece of identified DNA to discover phenotypes or activities in other protected databases, and *Completion Attacks* that use genotype imputation to uncover data that has been removed upon publication of a DNA sequence. To provide a complete overview of microbiome privacy risks, we detail each attack and then expand the categorization to include microbiome specific attacks ^1^.

*Identity Tracing With Metadata* reveals the identity of an anonymized DNA sample by using metadata such as age, pedigree information, geography, sex, ethnicity, and health condition. This attack is a concern with metagenomic comparative analysis as case and control group membership is determined by considering metadata.

*Genealogical Triangulation* uses genetic genealogy databases which link genealogical information, such as surname, with genetic material to allow an individual to recover ancestral information from his/her own DNA. This attack should not be a concern with microbiome data as microbiome inheritance has not been fully determined.

The microbiome presents three different methods for triangulation of a sample’s identity which we term *Location Triangulation*, *Behavior Tracing*, and *Rare Disease OTU*. As evidence of the first, a recent study detailed the similarity between individuals that occupy the same dwelling [14]. Therefore, an attacker may be able to reveal the identity of an individual microbiome sample by computing similarity with a sample taken from a specific location.

Further, *Behavior Tracing* could be used to identify a microbiome. The oral microbiota of romantic partners is more similar than other individuals and it is possible to measure how long the similarity between kissing partners is maintained [13]. An attack could be mounted using the phylogenetic or feature-level distance between a known person and the sample from a suspected romantic partner.

*Rare Disease Feature Tracing* takes advantage of attributes of public health disease tracking and microbial disease infections. Some infections, such as antibiotic-resistant cases, are recorded by state health departments and a single microbiome feature could correspond to those infections. If an attacker is able to observe the known microbiome feature of individual in a public health database and use it to link between another dataset, this will reveal any corresponding sensitive attribute.

*Identity Tracing by Phenotypic Prediction* involves predicting phenotypic information from genotypic information and then using that to match to an individual. Phenotypic prediction with human DNA is quite difficult given that predictions are not currently robust for unique identifiers in the population. For identifiers such as height, weight, and age, the effectiveness of this attack is likely to be low with microbiome data.

*Identity Tracing by Side Channel Leaks* is possible when an identifier is apparent from the dataset entries either by data preparation techniques or data-id assignment. One example is that Personal Genome Project sequencing files which by default were named with patient first and last name included. This attack is a concern with microbiome sequencing as well given that file uploading of the Personal Genome Project is similar for microbiome results.

*Attribute Disclosure With N=1* entails an attacker associating an individual’s identity to a piece of DNA and that piece of DNA to a sensitive attribute, such as an element in a database of drug users. For microbiome data, the forensic identification and the metagenomic codes techniques could be used by an attacker to successfully query a dataset with a sensitive attribute.

*Attribute Disclosure from Summary Statistics* uses genetic information of one victim and published summary statistics from a case/control study to determine if the victim’s DNA is biased towards the distribution of either the case or control group. If group membership can be determined, then the criteria to split groups (such as disease status) is revealed to the attacker. Linkage disequilibrium, or the probability that portions of DNA are more likely to be inherited together than others, provides a mechanism to increase the power of the attack. Further, genealogical information can be used to accomplish attribute disclosure.

While the authors cite Attribute Disclosure from Summary Statistics as an attack possible with all ‘-omic’ data, linkage disequilibrium and genealogical triangulation are not applicable to microbiome sequencing. The release of summary statistics may be used to determine if a metagenomic code for an individual is present in a case/control group, but the probability of this attack needs be determined.

*Completion Attacks* reveal portions of a DNA sample that are not released publicly by using linkage disequilibrium to uncover the hidden SNPs. Genealogical information, such as a pedigree and the SNPs of relatives, can also be used in genotype imputation. For metagenomic data, a cohabitation mapping of individuals from the same household to distinct features could be used to mount a completion attack.

## 3 System and Methods

In this section we first describe garbled circuits, which is the method we use for implementing secure metagenomic analyses. We then detail our system including participants, threat model, and approaches in the design space for privacy-preserving analysis.

### 3.1 Garbled Circuits

Two parties, one holding input *x* and another holding input *y*, wish to compute a public function over their inputs *F* (*x, y*) without revealing anything besides the output. The parties could provide their inputs to a trusted third-party to compute the function and reveal the output to each party, but modern cryptography offers a mechanism to run a protocol between only the two parties while achieving the desired functionality. The idea is to represent the function as a Boolean circuit over the inputs from both parties and use encryption to hide the input of each party during evaluation.

Secure function evaluation with garbled circuits occurs over several steps. First, Party 1 and Party 2 agree on a function (or, equivalently, a Boolean circuit) to compute over their inputs. Party 1 (the circuit “garbler”) constructs a garbled version of this circuit where each wire is associated with two “wire labels”, one of which is associated with the 0-bit and the other with the 1-bit on that wire. Each gate in the circuit is encrypted in such a way that one evaluating the garbled circuit can only derive one of the two output wire labels given one input wire label for each input of that gate. Party 1 sends this garbled circuit, along with the set of input wire labels associated with its input, to Party 2 (the circuit “evaluator”). The parties then run what is called an *oblivious transfer* protocol for Party 2 to receive the wire labels associated with its input. *Oblivious transfer* allows for a chooser, holding a 1-bit or 0-bit, to receive a message from a sender, holding two messages {M_0_, M_1_}, corresponding to its selected bit without revealing the bit to the sender as well as learning nothing about the other message held by the sender. Given the input wire labels for Party 1 and its own input, Party 2 can now evaluate the garbled circuit and learn the output, *without learning anything else about the circuit evaluation*.

The first practical implementation of garbled circuits was presented in 2004 [17]. Since then, the community has continued to develop implementations and provides efficient solutions for adversary levels including: (1) *semi-honest*, where an adversary follows the protocol but tries to infer the private input of the other party through analyzing the protocol transcript; (2) *covert*, in which an adversary attempts to cheat and is discovered with a defined probability [1]; and (3) *malicious*, where the adversary is allowed to deviate arbitrarily from the protocol description but should still not learn anything besides the function output.

### 3.2 System Participants

We consider the case in which parties that are located in two policy-domains want to perform metagenomic analyses over shared data. Examples of policy-domains include countries with differing privacy laws or institutions (universities, companies) that stipulate different data disclosure procedures.

For *i* ∈ 1, 2, denoting *P D_i_* as a Policy Domain, *R*_*i*_ as a researcher in Policy Domain *i*, *D*_*i*_ as the data from *R*_*i*_, *F* as the set of functions that a set of *R*_*i*_s would like to compute we consider the following setting:

> *R*_1_ and *R*_2_ would like to compute *F* over combined *D*_1_ and *D*_2_ but cannot do so by broadcasting the data as either *P D*_1_ or *P D*_2_ does not allow for public release or reception of individual-level microbiome data. We set *|i|* =2 but this setting could be generalized to any *i*.

Policy domains naturally arise due to differences in privacy laws. For example, studies currently funded by the NIH are required to release non-human genomic sequences including human microbiome data (http://gds.nih.gov/PDF/NIH_GDS_Policy.pdf). In contrast, the European General Data Protection Regulation, which is currently in draft form, lists biometric data and “any ‘data concerning health’ means any personal data which relates to the physical or mental health of an individual, or to the provision of health services to the individual” as protected information that is not to be released publicly (http://www.europarl.europa.eu/sides/getDoc.do?pubRef=-//EP//TEXT+TA+P7-TA-2014-0212+0+DOC+XML+V0//EN). Therefore, researchers in the US and EU may encounter different policies for data release but still have an interest in computing metagenomic analyses over shared data. Also, given the results published by Fransoza et al., some institutions may re-evaluate microbiome data release policies.

**Threat Model.** We consider a semi-honest adversary R1 who has a microbiome sample from a victim mixed with other samples. R2 is examining an association for a specific trait and would like to expand her study to use samples held by R1. R1 wants to determine if the victim is in a dataset of R2 and to learn a sensitive attribute of the victim such as disease status. We allow R1 to analyze the transcript and output of a set of metagenomic analyses that R1 and R2 agree to run.

Through using a garbled circuit implementation of metagenomic analyses, R2 will be able to keep the vector of microbiome features for any sample private, learn the outputs of functions that she would like to learn over the shared data, and prevent R1 from completing an *Identity Tracing* or *Attribute Disclosure with N=1* attack. As stated earlier, it is not clear if an *Attribute Disclosure from Summary Statistics* attack is a concern for metagenomic data. Finally, we defend against a *Forensic Identification* attack by restricting the set of functions so that R2 can prevent R1 from computing a distance-metric between any given pair of samples.

### 3.3 Solution Design Approaches

We consider different approaches to allow two parties to compute analyses over data which each must keep confidential.

**Access Control plus Trusted Third Party.** In the US, the NIH has recognized re-identification through publicly-posted genomic data as a realistic threat. Therefore, policy allows for publication of summary statistics and transfer of individual level sequencing data through access control using the Database for Genotypes and Phenotypes [16]. Once a researcher receives permission to access data, she is provided the data in an encrypted form along with a key to decrypt the data and operate over the data in the clear on her machine. We look to remove the need for access control by implementing the queries that a researcher would like to run without revealing the data directly.

**Differential Privacy.** While this approach provides provable privacy-guarantees, the introduction of statistical noise has not gained traction in the computational biology research community. Also, recent work showed that learning warfarin dosage models on differentially private datasets introduces enough noise that the dosage recommendation could be fatal to patients [9].

**Secure Multiparty Computation.** An alternative solution which we undertake, is using secure computation to perform metagenomic analyses. Other researchers have presented SMC for computing secure genome-wide association studies using secret-sharing, but that particular approach requires 3 parties and the use of third parties for computing tasks [11]. We address the feasibility of using garbled circuits to implement metagenomic analyses in terms of running time, network traffic, and accuracy. We believe that garbled circuits is the best approach for this scenario as it allows for direct communication between two parties and models the real-world setting well. Further, garbled circuits can handle a variety of adversaries beyond the semi-honest one that we consider in this work.

## 4 Implementation

In this section we describe how we implemented metagenomic analyses in garbled circuits and detail an evaluation of our system.

### 4.1 Metagenomics Using Garbled Circuits

We used an open-source secure computation library and note assumptions we make in handling our OTU count data. To implement each statistic, we focused on making the computations run in a feasible amount of time on real datasets.

**ObliVM.** ObliVM is a framework for secure computation including garbled circuits with a semi-honest adversary [15]. ObliVM allows for a user to write a function in Java for two parties to compute then compiles and evaluates the garbled circuit representation of that function. We implemented all metagenomic tests as Java packages then compiled and ran each with ObliVM. Our initial work on *χ*^2^-test was based on a *χ*^2^-test implementation using SNP data from https://github.com/wangxiao1254/idash_competition.

**Metagenomic Analysis Assumptions.** For this paper, we perform all analyses at the species level in secure computation. As detailed in Supplementary Note Section 1, OTUs are generated from direct pairwise comparison of sequencing reads. This is a compute-intensive process when performed on clear text. We do not attempt it in SMC for this work and assume each party performs this operation locally. We assume that each party will annotate each resulting OTU by matching to a common reference database, previously agreed upon by both parties (note that this reference database is orthogonal to sample-specific sequencing results obtained by each party). For illustration we assume that the agreed upon reference database yields annotation at the microbial species level. We also assume that parties can split data into case and control groups based on an agreed upon phenotype.

**Design approaches.** We took several approaches to implement each statistic. Since the metagenomic datasets we examined are at least 80% sparse and this trend is expected with OTU counts, we use sparse matrix computation techniques to make garbled circuits feasible [19]. For the sparse implementation, we provide only the non-zero elements for each feature from the dataset held by each party. We also operate per feature of the matrix as each OTU can be processed independently. These approaches allow us to amortize the overhead costs of evaluating the statistic for each feature by re-using the same circuit while changing the inputs.

In a faster approach, we use local pre-computation to reduce the number of operations performed in secure computation. For instance, the contingency table counts are computed locally for the *χ*^2^ test and odds ratio then the local results from each party are combined in the garbled circuit. To measure the impact of our design choices we implemented a naive algorithm for each statistic and compared results. We provide greater detail for each implementation of the *χ*^2^ test, odds ratio, Differential Abundance, and Alpha Diversity in the Supplementary Note, Section 3.

### 4.2 Evaluation

We evaluated our implementation using two Amazon EC2 

~~~
r3.2xLarge
~~~

 instances with 2.5GHz processors and 61 GB RAM running Amazon Linux AMI 2015.3. We measured the size of the circuit generated, running time, and network traffic between both parties for each metagenomic statistic and dataset. Circuit size serves as a useful comparison metric since it depends on the function and input sizes but is independent of hardware. Running time and network traffic are helpful in system-design decisions and benchmarking of deployments.

### 4.3 Datasets

We used OTU count data from the Personal Genome Project (PGP) [5], the Human Microbiome Project (HMP) [23], and the Global Enterics Multi-Center Study (MSD) [20]. The MSD data was retrieved from ftp://ftp.cbcb.umd.edu/pub/data/GEMS/MSD1000.biom while the PGP and HMP datasets are from https://github.com/biocore/American-Gut/tree/master/data [4]. After aggregating to species and removing features which hold all zeros for either the case or the control group, the PGP contains 168 samples and 277 microbiome features, the HMP has 694 samples and 97 features, and the MSD dataset consists of 992 samples and 754 features. Supplementary Table 2 summarizes the size and sparsity of each dataset.

### 4.4 Efficiency of Secure Computation

**Circuit Size.** Figure 1 shows the circuit size per feature for each experiment. Using pre-computation, the complexity of the equation to calculate each statistic determines the circuit size. This explains the circuit sizes for odds ratio and *χ*^2^ test as compared to Differential Abundance. For Alpha Diversity, all rows and columns are pre-processed with only the two sample t-test computed in the garbled circuit. With the sparse implementation, the complexity of the test along with the number of non-zero elements in the dataset directly affects circuit size.

**Figure 1:**
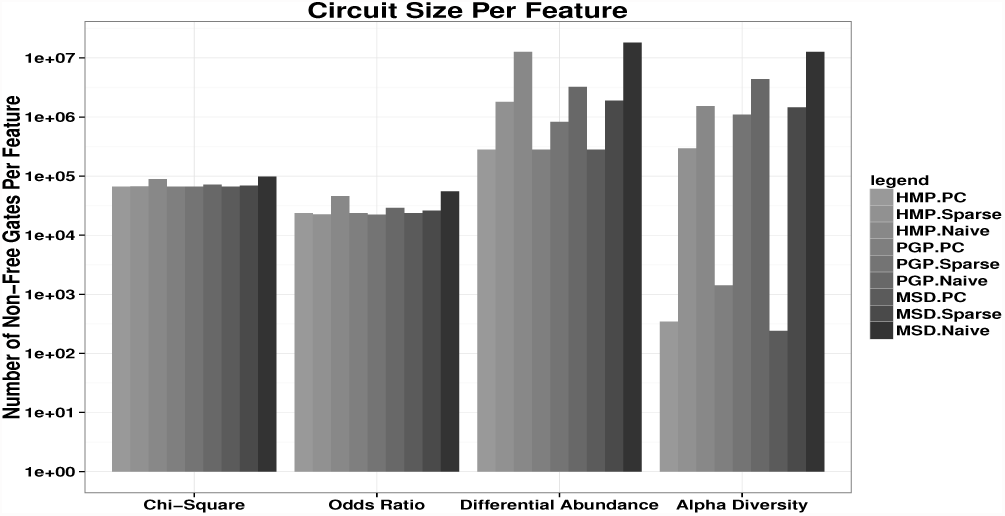
PC stands for Pre-Compute. Circuit size for each implementation and dataset. The features for Alpha Diversity is the number of samples. The differences in Alpha Diversity between datasets is explained by the number of samples for PGP (168) is much lower than that of HMP (694) and MSD (992).

**Running time.** For the sparse implementation, the running time was proportional to the size and number of non-zero elements in each dataset. For pre-computation, Alpha Diversity was affected by the number of samples in each dataset. The running time for the *χ*^2^ test, odds ratio, and Differential Abundance were proportional to the number of features (rows) processed. Figure 2 summarizes the effects of input size and algorithm complexity on running time.

**Figure 2:**
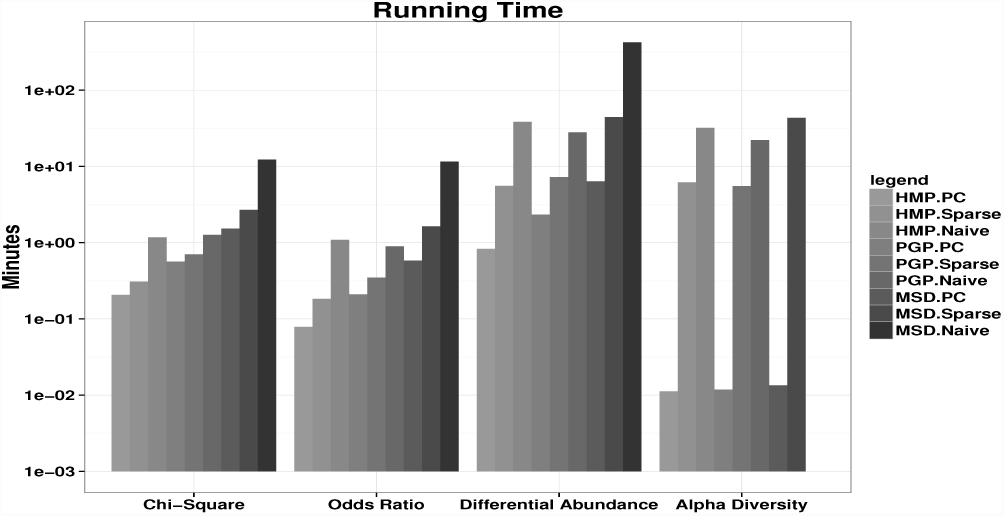
PC stands for Pre-Compute. Running time for each statistic and each dataset in minutes. In each statistic, the number of arithmetic operations determined the running time. The size of the dataset along with sparsity contributed to running time for the sparse implementations. Alpha Diversity MSD Naive did not run to completion on the EC2 instance size due to insufficient memory. Based on the circuit size and the number of gates processed per second for other statistics, we estimate the running time to be 378 minutes.

**Network traffic.** Supplementary Table 3 shows the network traffic for each experiment. The increase in network traffic between the pre-computation and sparse implementations is more significant than the differences in running times of those approaches. We believe that the network traffic for the pre-compute implementation is quite good for the security guarantees provided with using garbled circuits while the sparse approach presents an acceptable tradeoff depending on the network resources available.

### 4.5 Accuracy

We compared the accuracy of our implementation results to computing the statistic using standard R libraries. Table 1 lists the accuracy of results for the *χ*^2^ statistic along with p-values, odds ratio, and Differential Abundance ttest results. The differences in our results between the R values appear to be the result of floating-point rounding errors.

**Table 1:** Comparison of results generated using the R chisq.test{stats}, odds.ratio{abd}, t.test stats, and diversity {vegan} against our implementation in **ObliVM-GC** for the *χ*^2^ test, odds ratio, Differential Abundance, and Alpha Diversity. We use Normalized Mean Squared Error: ||*x* − *y*||^2^/||*x*||^2^ with *x* as the value output by R and *y* the value from our implemenation. For comparing p-values, we use the log10 p-value and exclude any exact matches (since log_10_(0) = -Inf in R) while computing the mean.

|  | PGP | HMP | MSD |
| --- | --- | --- | --- |
| Chi-Square statistic | 7.84e-07 | 7.48e-06 | 7.02e-08 |
| Chi-Square p-value | 2.00e-07 | 2.14e-06 | 9.72e-08 |
| odds ratio | 1.60e-13 | 5.42e-13 | 2.44e-13 |
| Differential Abundance<br>t-statistic | 0.023 | 0.0017 | 0.0012 |
| Differential Abundance<br>degrees of freedom | 2.7e-4 | 2.5e-4 | 0.0028 |
| Differential Abundance<br>p-value | .0024 | 0.0026 | 0.0011 |
| Alpha Diversity<br>t-statistic | 0.0038 | 0.017 | 0.0049 |
| Alpha Diversity<br>degrees of freedom | 1.48e-05 | 9.7e-4 | 2.2e-4 |
| Alpha Diversity<br>p-value | 0.0088 | 0.044 | 0.014 |

We investigated if our implementation yielded any false positives and false negatives with the results from R acting as ground truth. For the p-values of Differential Abundance in PGP, HMP, and MSD datasets we found no false positives or false negatives for a significance level of 0.05.

### 4.6 Significant Features Discovered Through Data-Sharing

Researchers in different policy domains may be forced to compute analyses on partial data. We measured the effect of using our implementation for data-sharing between policy domains. The MSD dataset provides a means to simulate secure computation of microbiome analyses between different countries. The data was gathered from Kenya, The Gambia, Bangladesh, and Mali. We simulate each country performing secure Differential Abundance pair-wise with the other countries. We observed that sharing data resulted in a substantial increase (at minimum a 98% increase) in the number of species found to be differentially abundant between case and control groups. Table 2 summarizes the results.

**Table 2:**
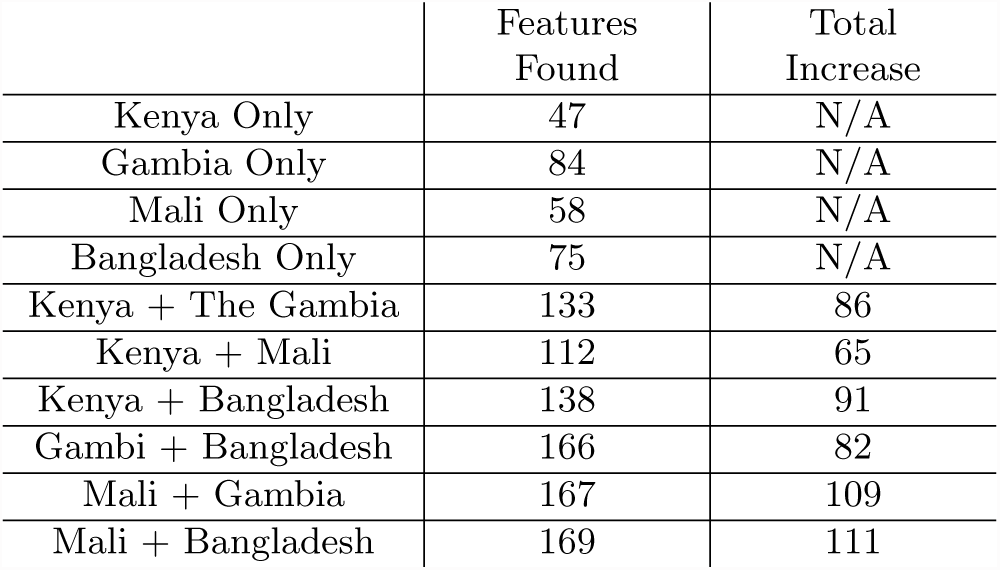
Significant Features Found From Sharing Data Between Each Country. When computing data with another policy domain, each country saw an increase in the number of features detected to be significantly different between case and control groups.

### 4.7 Metagenomic Codes

We also evaluated our implementation on the genetic marker data that showed the greatest identification power in the metagenomic codes analysis [8]. The data is also from the HMP and consists of a total of 85 samples and 221,111 features. Due to the large number of features and sparsity of the data, we implemented a filtering garbled circuit in which we first return a vector to each party denoting if a given feature meets a presence cutoff and then have each party input those features into our existing implementations to compute the statistical test. For *χ*^2^, the 1,729,851,751 gate circuit is evaluated in 67.4 minutes, with 51,926.35 MB sent to the evaluator, and 1,642.53 MB sent to generator. For odds ratio, the 632,918,505 gate circuit is evaluated in 33.18 minutes, with 1,642.29 MB sent to the evaluator, and 20,542.84 MB sent to generator. This results shows that the secure comparative analyses we would like to perform are possible given the legitimate concerns raised by Franzosa et al.

## 5 Discussion

In this section we describe related work and provide a context for our contribution. We also discuss a use case for our solution in building datasets and finally present conclusions we formed during the course of our work.

### 5.1 Related Work

As we are the first, to our knowledge, to approach secure microbiome analysis, we review related work on privacy-preserving operations over human DNA.

**Secure DNA Sequence Matching and Searching.** Comparing two DNA segments is essential to genome alignment and identifying the presence of a disease causing mutation. One approach is to use an oblivious finite state machine for privacy-preserving approximate string matching [21]. Also, the garbled circuits technique can be used to implement secure text processing and has applications to DNA sequence comparison [12]. FlexSC, a widely-used garbled circuits implementation which ObliVM-GC is derived from, was benchmarked by computing Levenstein distance and the Smith-Waterman algorithm between private strings held by two parties [10].

**Privacy-Preserving Genome-Wide Association Studies.** Prior work has shown that secure computation between two institutions on biomedical data is possible by using a 3-party secret-sharing scheme [11]. The authors present an implementation of a *χ*^2^ test over SNP data using the Sharemind framework. Other researchers have presented a modification of functional encryption that enables a person to provide her genome and phenotype to a study but only for a restricted set of functions based on a policy parameter [18].

**Secure Genetic Testing.** For using sequencing results in the clinical realm, paternity determination and patient-matching is possible using private set intersection [3]. Also, it is feasible to utilize homomorphic encryption for implementing disease-risk calculation without revealing the value of any genomic variant [2].

### 5.2 Patient Pool

The recent announcement by 23andMe to begin drug development on its genome variant datasets highlights the value of biomarker data. We imagine a scenario where individuals can use our solution to create and manage datasets in order to charge drug developers to run analysis functions over the data. The companies will have to be non-colluding as otherwise all function results could be shared among companies. The current regulatory process for drug development allows a mechanism to enforce this constraint.

The patient pool can be paid to compute a function to over its data and sign the output either with one public key for the pool or using a multiple signature scheme between all pool participants. Upon requesting drug trial permission in the US, a company is required to hand over all data from research, which in this case would include the output of the patient pool analysis and signatures over those results. The FDA could verify the signatures to enforce non-collusion between companies. We believe this provides a mechanism to create high-quality datasets that are accessible to a variety of companies and ensure patients are compensated for their efforts.

Formally, we denote *P* as a patient, *D* as the biomarker data for patient *P*, *M* as a patient pool manager, *R* as a researcher, *F* as the function that *R* would like to compute by combining its own data with the *D*s held by *M*, and *A* as a regulatory authority.

*M* is trusted not to reveal the data of any *P* in the pool and is semi-honest with regard to the interaction with *R*. *R* is semi-honest and will not attempt to learn anything beyond what can be inferred by the output of *F* and inspection of the protocol transcript. *A* is a trusted authority that can verify a signature over the output of *F* from a pool managed by *M*. We assume the existence of a Public Key Infrastructure which links the identity of *M* to a given public-key. In practice, 23andMe or uBiome could act as an *M*, *R* could be a pharmaceutical development company, and *A* would the FDA or European Medicines Agency. Supplementary Figure 2 details the specific interactions between each system participant.

### 5.3 Conclusions

In this paper we have provided a categorization of privacy issues in the analysis of metagenomic sequencing. In addition, we have shown that it is possible to perform metagenomic analyses in a secure computation framework. Our implementation made use of pre-computation steps to minimize the number of operations performed in secure computation making the use of garbled circuits feasible. We also implemented sparse-matrix methods for each statistic. We took this step in order to prove the applicability of this solution for other analyses when the data itself acts as sufficient statistics, such as for the Wilcoxon rank-sum test.

We believe the patient pool idea will benefit patient groups, specifically those suffering from rare diseases or those with insufficient data in existing repositories for association studies. Also, this could enable faster and higher quality drug development as drug-companies will have a richer set of data to examine.

While the storage and sharing of medical data is ultimately a policy matter, providing a technical solution is useful to forming good policy. We believe that given the time costs associated with re-consenting patients to release data to another researcher or creating a legal contract stipulating a data receiver’s responsibility, that the running times we presented for metagenomic analyses are a reasonable tradeoff.

DNA sequencing technologies are entering a period of unprecedented applicability in clinical and medical settings with a concomitant need for regulatory oversight over each individual’s sequencing data. We believe that addressing privacy concerns through computational frameworks similar to those used in this paper is paramount for patients while allowing researchers to have access to the largest and most descriptive datasets possible. We expect that secure computation and storage of DNA sequencing data, both the individual’s DNA and their metagenomic DNA, will play an increasingly important role in the biomedical research and clinical practice landscape.

1 We use the names for each attack as introduced by Erlich and Narayanan.

## 6 Acknowledgements

**Funding:** This work was partially supported by the National Institutes of Health R01HG005220 to H.C.B and J.W.; the US National Science Foundation Graduate Research Fellowship DGE0750616 to J.N.P; and the US National Science Foundation 1111599 to X.S.W, 1314857 to X.S.W.

